# Fungal symbionts of Arctic zooplankton induce mortality and reduce swimming activity

**DOI:** 10.1101/2020.09.24.312272

**Authors:** BT Hassett, E Ershova, J Berge

**Author notes:** **Corresponding author:** Brandon T. Hassett.

## Abstract

Zooplankton are key intermediaries between microbes and higher trophic levels. Elevated temperatures in the Arctic are expected to increase metabolic rates, which in turn will result in increased rates of zooplankton production and foraging. We postulate that this may also be tied to elevated incidence of fungi parasitism in key zooplankton species. We present data of *in-situ* fungal parasitism in the key Arctic zooplankton species complex *Calanus* spp. We found that >10% of *Calanus* specimens were internally parasitized by fungi that had 100% molecular identity to *Penicillium chrysogenum*. Experimental studies with two strains of *P. chrysogenum* and *Calanus* spp. revealed an increased mortality effect suggesting that fungi decrease the survival of overwintering zooplankton, which may have cascading effects on higher trophic levels that feed on *Calanus*. Moreover, *Calanus* demonstrated reduced swimming activity in the presence of *P. chyrsogenum*, indicating that fungi could affect predator avoidance and vertical migration patterns of zooplankton. Semi-thin sectioning of live zooplankton incubated with *P. chrysogenum* revealed the presence of hyphae throughout the body, underscoring the parasitic nature of *P. chrysogenum* in *Calanus*. We hypothesize that as the Arctic warms, the balance between host and pathogen could shift in favor of pathogen reproduction, resulting in a greater effect of fungal pathogens on zooplankton.

## Introduction

The Arctic marine ecosystem is shaped by a strong seasonality of light and nutrient fluxes that results in distinct seasonal peaks of primary production (Gradinger 2009; Arrigo et al. 2012). The link between primary production and higher trophic levels can be particularly strong in the Arctic (Chassot et al. 2010), where ecosystem complexity is characterized by seasonally and regionally occurring short food webs (Iken et al. 2010; Darnis et al. 2012). Key zooplankton species, such as *Calanus* spp., capitalize on the brief sea ice and phytoplankton blooms, reducing their metabolism during the winter months and relying on accumulated lipid stores to survive the low-productive period. Major shifts in the overwintering success of zooplankton, therefore, presumably will have cascading effects on higher-trophic levels, such as pelagic fish and the ice-associated polar cod (*Boreogadus saida*) that feeds on calanoid zooplankton (Renaud et al. 2012; Rand et al. 2013). Polar cod, in turn, is a major prey item for birds and many ice-associated seals(Bluhm and Gradinger 2008).

Marine fungi are globally distributed (Tisthammer et al. 2016; Hassett et al. 2019b; Morales et al. 2019) and comprise comparable quantities of biomass (~1 mg Cm^-3^) as other abundant marine taxa (Damare and Raghukumar 2008; Gutiérrez et al. 2011; Bochdansky et al. 2017; Hassett et al. 2019a). Increasing quantities of diverse lines of evidence underscore that fungi are nearly ubiquitous in the marine realm (Burgaud et al. 2009; Edgcomb et al. 2011; Orsi et al. 2013; Taylor and Cunliffe 2016; Li et al. 2018; Garvetto et al. 2019; Pang et al. 2019), including throughout the Arctic (Sparrow 1973; Pang et al. 2008; Terrado et al. 2011; Zhang et al. 2015; Hassett and Gradinger 2016; Hassett et al. 2019a). Fungi are diverse (Kagami et al. 2014) and many marine taxa represent novel, ecologically uncharacterized lineages(Richards et al. 2012; Ettinger and Eisen 2019), including in the Arctic Ocean (Comeau et al. 2016; Hassett et al. 2017). This hyper-diversity within the Fungi correspond to varied life history strategies, with an associated broad spectrum of morphology circumscribing intracellular organisms (Quandt et al. 2017), ~2 μm reproductive zoospores (Scholz et al. 2016), and extensively branched hyphae (Gutiérrez et al. 2011).

As the Arctic continues to warm, large scale physical phenomena, such as the loss of sea ice, are driving shifts in biological communities and resulting in the northward migration of boreal taxa (Wesławski et al. 2011; Fossheim et al. 2015). The effects of these physical changes are pronounced on microbial communities (Comeau et al. 2011) that can rapidly shift into new equilibrium states based on relatively short doubling times. In addition to their biogeochemical cycling abilities, microbes are also parasitic, whose ability to cause disease is partially governed by environmental factors (Scholthof 2007; James et al. 2015). Fungi are parasites of marine mammals (Higgins 2000; Miller et al. 2002; Rosenberg et al. 2016), crab eggs (Shields 1990), sea fans (Kim et al. 2006), sponges (Gao et al. 2008), and zooplankton (Seki and Fulton 1969). In zooplankton, fungal parasitic activity is governed by temperature and rates of host foraging (Shocket et al. 2018). This is of particular interest, as the Arctic is rapidly warming and fungi have been detected in the gut content of high-latitude marine zooplankton, suggesting active feeding (Cleary et al. 2016, 2017; Yeh et al. 2020). Though fungi are regularly detected in association with metazoans, the nature of this co-occurrence (e.g. mutualism, commensalism, parasitism) remains to be elucidated. Mutualistic anaerobic fungi within the Neocallimastigomycota inhabit the guts of mammalian herbivores (Gruninger et al. 2014) and have been reported from marine systems (Hassett and Gradinger 2016; Picard 2017). Moreover, fungi are known pathogens of select copepod species in marine ecosystems (Seki and Fulton 1969). These variable symbiotic relationships underscore the importance of understanding interactions beyond co-occurrence to understand the ecology.

The majority of high-latitude marine fungal studies have strategically moved to establish baselines of species richness and diversity through culturing and molecular surveys. Though it is presumed that molecular detection and cultivation of fungal species correspond to presence and *in situ* activity, experimental evidence demonstrating ecological roles remain to be generated, as in other marine environments (Cunliffe et al. 2017). Recent studies have suggested that *Calanus* spp. play a significant ecological role also during the dark polar night(Berge et al. 2015), but have also documented mortality rates during the polar night (Daase et al. 2014). We assessed the *in situ* incidence of fungal symbionts in *Calanus* spp., as well as conducting *in vitro* experiments to assess their effect on mortality and behavior. In order to link the studies to over-wintering survival, all *in situ* sampling were carried out during the arctic polar night period.

## Methods

### Overview

Zooplankton sampling was conducted in two high Arctic locations in the Svalbard archipelago (Kongsfjorden and Billefjorden) during the polar night in January 2020, with additional sampling for zooplankton conducted in Ramfjorden (northern Norway) during February 2020. Zooplankton in both locations were sampled using 180μm WP2 nets (Hydro-Bios, Table 1). The dominant species *Calanus* spp. (a mix of *C. glacialis* and *C. finmarchicus*, stage CV/adult females) were live-sorted and incubated in the dark at 4 °C until processing (within 48 hours of collection).

**Table 1.** Summary of fungal survey and experiments. *In situ* incidence summarizes the number of fungi screened by location for the presence of fungal symbionts. Zooplankton mortality shows the percent of *Calanus* spp. that died during incubation experiments. Swim counts is the average number of vertical migrations over 26.5 hours with standard deviations. ZP1 and ZP2 are two strains of *Penicillium chrysogenum* cultured during *in situ* studies and introduced during experiments.

| ***In situ* incidence** |  |  |
| --- | --- | --- |
|  | **# *Calanus* spp.** | **# *Calanus* spp. with fungi** |
| Billefjorden | 16 | 0 |
| Kongsfjorden | 93 | 10 |
| Ramfjorden | 71 | 12 |
| **Zooplankton mortality** |  |  |
| **Control** | **ZP1** | **ZP2** |
| 35% | 32% | 84% |
| **Swim counts** |  |  |
| **Control** | **ZP1** | **ZP2** |
| 487 (450) | 267 (307) | 148 (191) |

At the Svalbard locations, the pre-sorted *Calanus* spp. were explored for the presence of endogenous fungal symbionts onboard the sampling vessel (FF *Helmer Hanssen* on Svalbard). Live and dead *Calanus* (87 total, 16 from Billefjorden and 71 from Kongsfjorden) were surface sterilized in absolute ethanol for 30 seconds, rinsed in autoclaved seawater, and subsequently incubated for one week at room temperature in sterile Petri dishes in moist chambers onboard the ship. After this first survey, individuals that grew fungi were transferred to PmTG agar media (Barr 1986) that was amended with seawater, as well as streptomycin and penicillin g antibiotics. Fungal isolates were subcultured, until an axenic culture was established. Though means were taken to eliminate contamination onboard the ship, restrictions governing open flames left the small possibility that zooplankton were contaminated by air-borne fungal spores. As a follow-up to this ship-based study, Ramfjorden *Calanus* spp. (93 individuals) were collected and subsequently processed as described above in a laminar flow hood under strict sterile conditions (bleach and ethanol sterilization, followed by UV treatment for 30 minutes). Glass slides were extensively flamed, as were forceps between each zooplankton picking event. To control for the ethanol surface sterilization efficacy, zooplankton were dragged and rolled extensively in a pure culture of our fungal isolate, until the exoskeleton was visibly covered in fungal spores. Both positive (incubated immediately after fungal coating) and negative (surface sterilized in absolute ethanol and rinsed in autoclaved seawater) controls were incubated in separate moist chambers, in tandem with our other zooplankton.

Pure cultures of our two fungi (strain ZP1 and ZP2) were DNA extracted using the DNeasy PowerMax Soil extraction kit, per manufacturer instructions. The fungal ITS locus was amplified using PCR with the ITS4-ITS5 primer set (White et al. 1990) and Sanger-sequenced bidirectionally by GeneWiz UK. Sequences were explored for the presence of ambiguous peaks, aligned in MEGA, and queried with NCBI’s BLAST to identify sequences with closest identity. High-quality sequences were deposited in NCBI under accessions MT218320-MT218321.

### Zooplankton Activity Chambers and Experimental Mortality

Suspension of fungal spores from strains ZP1 and ZP2 were made in autoclaved seawater and subsequently enumerated with a hemocytometer. A total of 30,000 fungal spores from each strain were separately added to five 250 mL culture flasks (for a total of 10 flasks containing fungi) full of 0.2μm-filtered seawater. Each flask contained five live individuals of adult female *Calanus* spp. Eight control bottles (no fungi) containing the same number of zooplankton were established and monitored in parallel with our experimental bottles. Bottles were incubated for five weeks at 10 °C. At the end of five weeks, mortality was recorded.

After the five week incubation, the remaining live copepods were placed into individual 10 mL glass tubes filled with filtered seawater and their activity level was monitored for 24 hours using Trikinetics Locomotor Activity Monitors (LAMs), which quantify vertical swimming activity as the number of laser beam breaks across the center of the tube, per unit time. A total of 30 copepods were incubated in this way (11 controls, 11 individuals co-incubated with ZP1, and 8 individuals co-incubated with strain ZP2). The LAMs were kept at 4 °C in the dark for the duration of the experiment.

### Semi-thin sectioning and light microscopy

Samples in alcohol were cut in two and fixed in 2.5% glutaraldehyde in PHEM buffer overnight. Samples were post-fixed in 1 % aqueous OsO4 and washed in PHEM buffer then water and subsequently stained in 2 % uranyl acetate before dehydration in a graded series of ethanol, followed by 2 x propylenoxide. The samples were incubated in 50:50 propylenoxide epoxy resin (Serva Agar 100 R1043, DDSA R1051, MNA R1081, DMP-30 R1065) overnight, followed by in pure resin with polymerization at +60°C for 48 hours. Semi-thin and semi-thin sections were cut on a Leica Ultracut UM7 ultramicrotome (Vienna, Austria) with a glass knife (semi-thin) and diatom knife (Diatome, Bijel, Switzerland). Semi-thin sections were stained with 1% toluidine blue and scanned on an Olympus VS120 Slide Scanner.

## Results

### Fungal incidence

Our broad survey of the presence of endogenous fungi in *Calanus* spp. from Svalbard during the polar night 2020 cruise revealed that 11% (10 ethanol-sterilized specimens) hosted endogenous fungi. All ten of these individuals (8 living specimens and 2 carcasses) were collected from Kongsfjorden (14% of local total). All fungi that grew in zooplankton were hyphal-forming with dark-green conidia. After culturing, DNA extraction, and BLAST queries, both isolates were allied to *Penicillium chrysogenum* (100% identity, 100% query coverage). None of the specimens from Billefjorden showed signs of hosting fungi.

Replication of this study in Ramfjorden under strict sterile conditions revealed that 13% (12 female *Calanus* spp.) hosted endogenous fungal symbionts. All fungi produced hyphae with dark-green spores, whose morphology closely resembled our sequenced isolates. Inspection of these zooplankton after one-week incubation at room temperature revealed extensive hyphae protruding from prosomite segments (Figure 1A, B), as well as extensively through the body of copepods (Figure 1C), eventually resulting in external sporulation (Figure 1D-F). Negative controls that were coated in spores and sterilized with absolute ethanol, did not yield any growth after one week. Positive controls showed extensive fungal growth.

**Figure 1.**
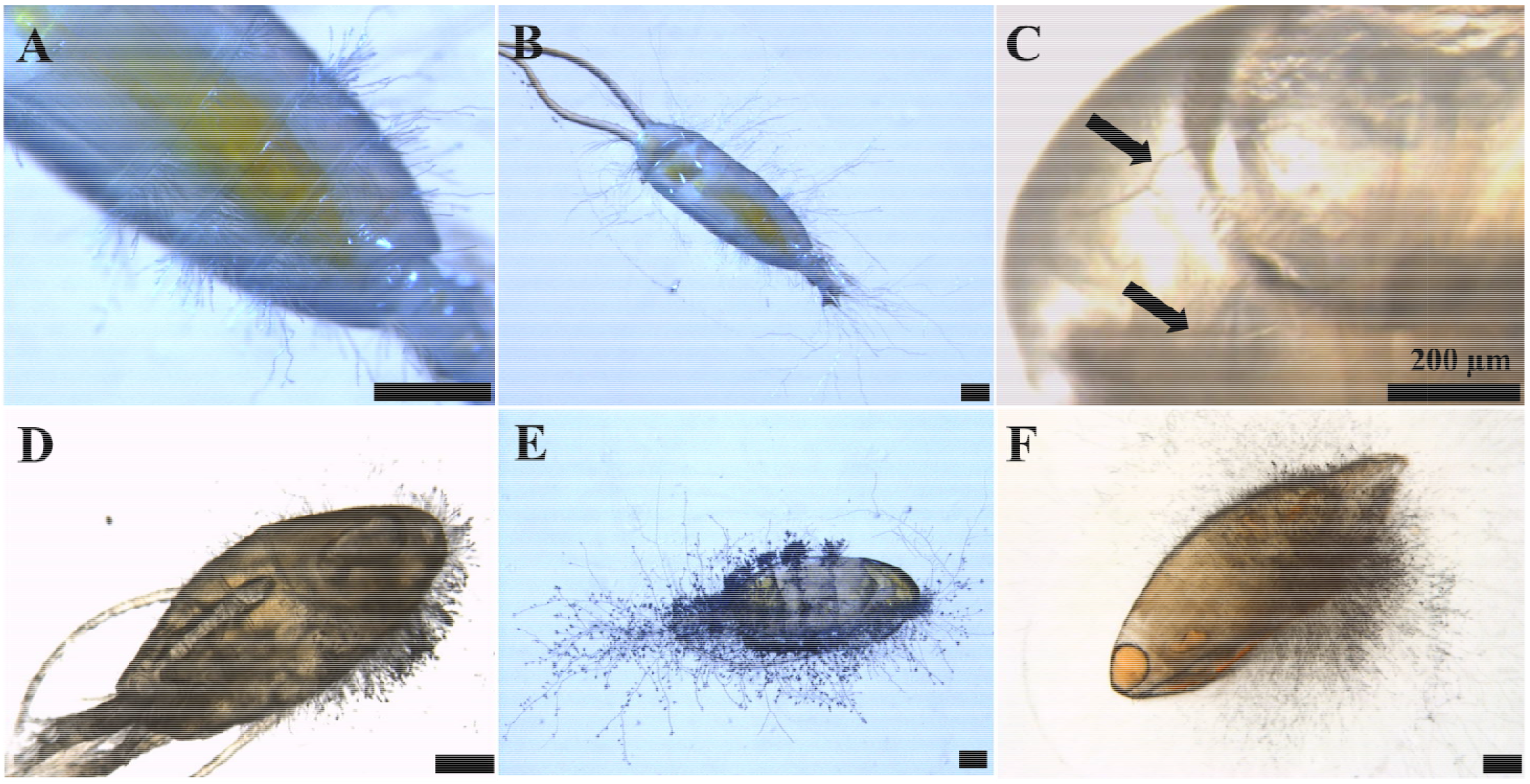
Overview of *Calanus* five days postmortem. A) Fungal hyphae protruding from prosomite segments. B) Overview of Figure A with extensive hyphae radiating from zooplankton body. C) Endogenous hyphal filaments at head of *Calanus* (arrow). D-F) Representative photos of mature fungal conididiophores sporulating externally on copepod body. Scale bar represents 500 μm, unless noted otherwise.

### Mortality and activity

After five weeks of live incubations, 65% of control zooplankton remained alive and active (Table 1). Despite the relatively high mortality in control bottles, experimental bottles containing *P. chrysogenum* had higher mortality than the control zooplankton incubated in the absence of fungi (p=0.26, two-tailed T-test). Specifically, we recorded a heightened mortality effect associated with the co-occurrence of Strain ZP2 (p=0.03, two-tailed T-test) than ZP1 (p=0.90, two tailed T-test). To follow-up on this perceived difference of strain-specific fungal activity on calanoid zooplankton, we monitored behavior over a 24-hour period in zooplankton that remained living after five weeks. During the course of the behavior experiment, several specimens died, leaving a total of 7 control *Calanus*, 9 ZP1 *Calanus*, and 5 ZP2 *Calanus*. We found that the activity level in the surviving zooplankton varied between treatments, with the control group containing the most active individuals. Specifically, control zooplankton had an average of 487 (± 450) swimming events per 26.5 hours; *Calanus* with ZP1 had an average of 267 (± 307); and *Calanus* with ZP2 had an average of 148 (± 191) swimming events over the same period, corresponding to an overall reduced swimming events between controls and *Calanus* co-incubated with *Penicillium* (p=O. 19, two-tailed T-test) with strain ZP2 having a more pronounced effect on behavior (p=0.10, two-tailed T-test) than ZP1 (p=0.29, two-tailed T-test). As parasitism is governed by both host and parasite genetics, we inherently assumed and applied unequal variances to our statistical analyses because we used two different fungal strains.

### Semi-thin sectioning and microscopy

After incubations, live zooplankton were selected for electron microscopy. Several of these live, swimming zooplankton appeared green under the dissecting scope. After embedding, sectioning, and counter staining, fungal-like structures were observed throughout the zooplankton sections, including long hyphal-like strands and transverse sections of hyphae. Specifically, zooplankton cells stained blue due to their DNA; however, transverse sections of hyphae generally do not contain nuclei, as nuclei congregate at the apical end of growing hyphal strands. As a result, transverse sections of hyphae appear more purple. These hyphal sections have variable diameter and appear throughout zooplankton body, seemingly not localizing at the central core, where lipid stores and the gut content is localized (Figure 2).

**Figure 2.**
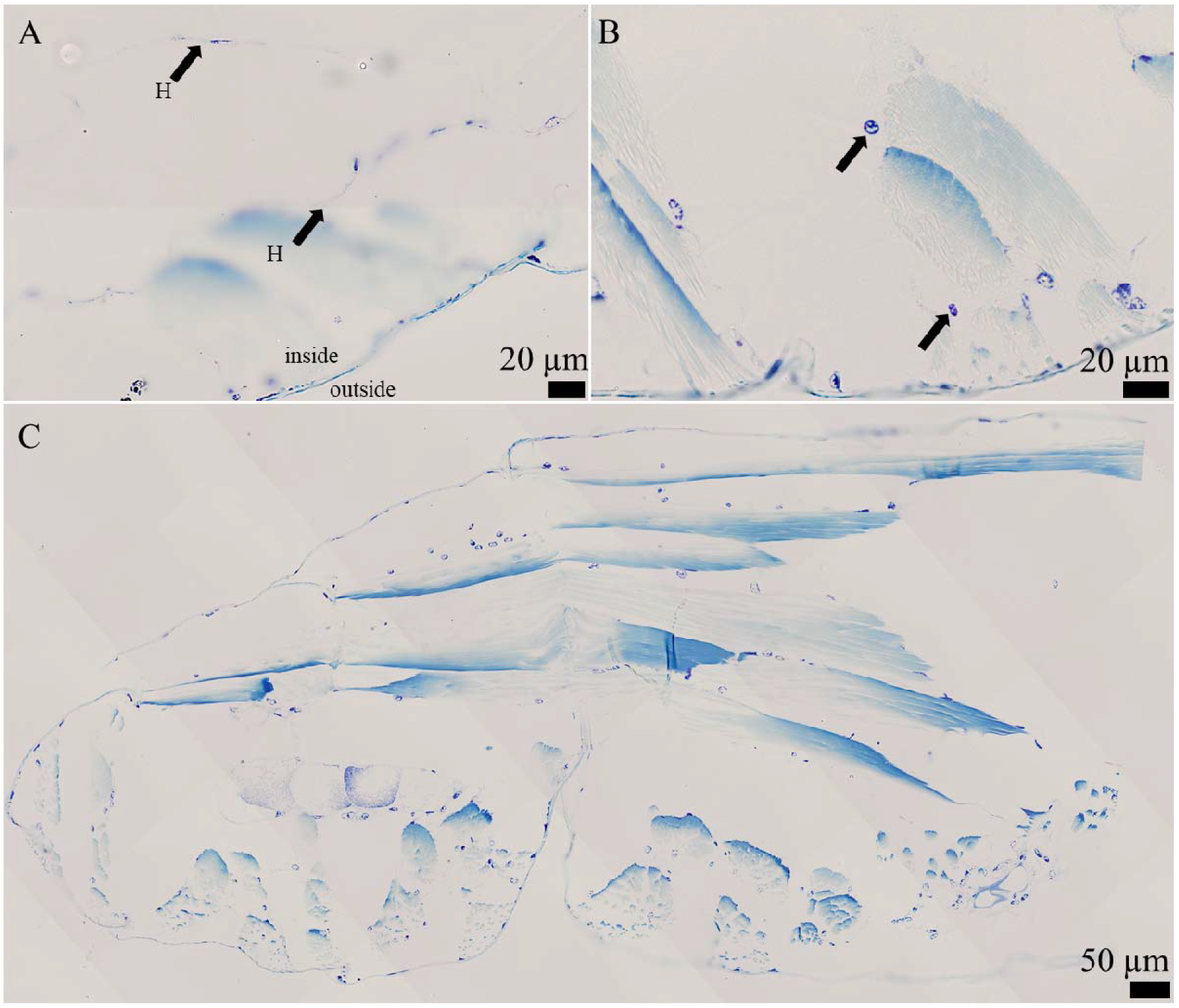
Semi-thin sections of *Calanus* zooplankton co-incubated with *Penicillium chrysogenum*. Living *Calanus* specimens were selected for sectioning. A) Hyphal filaments (H) on the inside of *Calanus*. B). Longitudinal sections of hyphal filaments (arrow) on the inside of zooplankton. C) Large overview of a zooplankton cross section.

## Discussion

Our analysis revealed surprisingly high (>10%) co-occurrence of endobiotic *Penicillium* species inside *Calanus* at two different locations in the Arctic. Hyphae were visibly protruding from the inside of carcasses, definitively proving the endobiotic nature of these fungi. *Penicillium* species are globally distributed (Hassett et al. 2019b), produce a suite of degradative enzymes (Park et al. 2019), are commonly found in seafood (Park et al. 2019), and can be abundant in the Arctic (Luo et al. 2020). Our mortality experiments indicate that the presence of select *P. chrysogenum* strains can differentially reduce survival in copepods during a time span of weeks, consistent with variable virulence among strains of the same fungal species (Kowalski and Cramer 2020). Non-consumptive mortality of zooplankton in the Arctic may peak during winter months, as evidenced by sediment traps, which found *Calanus* carcasses to be the dominant component of sinking carbon outside of the spring bloom (Sampei et al. 2012). Similarly, zooplankton surveys during January in the Svalbard area found that up to 95% of *Calanus* specimens found deeper than 300m, and up to 50% in the surface layers, were dead, with many of them still containing non-depleted lipid stores (Daase et al. 2014). The causes for these mortality events are unknown, but diseases and parasites are among the suspected culprits (Tang and Elliott 2013). Our experimental results provide circumstantial support for a fungi-induced mortality rate in over-wintering *Calanus* spp.

If fungal parasitism does not result in mortality, it is presumed that mounting a defense response comes at an increased metabolic cost, likely to the detriment of lipid storages that sustain life during the polar night (Berge et al. 2015). Symbiotic fungi can produce a suite of secondary metabolites (Gao et al. 2011; Nielsen et al. 2017) that might affect the behavior of zooplankton, as is reported for fungi in other invertebrates (Hughes et al. 2011; de Bekker et al. 2014). Our results demonstrate that both non-virulent and virulent strains of *P. chrysogenum* can reduce swimming activity in calanoid zooplankton. Although *Calanus* supposedly overwinter at depths below 100m, recent studies show that a large portion of the population remains in the surface layers (Kvile et al. 2019). We provide a possible explanation for the different behavior in individuals belonging to the same population and for why many of these zooplankton remain at depth: they are colonized by fungi that reduce their behavior. For zooplankton hosting *Penicillium* symbionts, irrespective of direct mortality, we hypothesize that these fungi degrade zooplankton carcasses, as other fungi do in freshwater ecosystems (Tang et al. 2006; Tang and Elliott 2013), mediating the amount and quality of carbon that will eventually reach the sea floor and remain in the water column. The disease incidence and effect of other parasites on zooplankton remains to be elucidated.

As temperatures increase in the Arctic, the environmental factors that regulate disease, including temperature, could shift to favor pathogen reproduction (Vaumourin and Laine 2018) with unknown consequences on host populations or higher trophic levels that feed on those hosts. Environmental forces, such as advection of pathogens into conducive areas, may amplify effects of pathogens on animals, as anecdotally supported by the relatively high infection rate in the advection-dominated Kongsfjorden, but not the oceanographically isolated Billefjord (Berge et al. 2014). The species composition is also more skewed towards the Arctic *Calanus glacialis* vs the Atlantic *C. finmarchicus* in Billefjorden compared to both Kongsfjorden and Ramfjorden (Berge et al. 2014). However, our study was not designed to determine a more precise species affinity of *P. chrysogenum* towards *C. finmarchicus* (vs *C. glacialis)*. As zooplankton metabolism rates will likely increase with increasing Arctic temperature (Alcaraz et al. 2013), the positive relationship between temperature and zooplankton foraging rates, as well as zooplankton foraging rates and fungal disease prevalence (Shocket et al. 2018) is particularly topical for Arctic marine ecosystems. Ultimately, these data demonstrated that *Penicillium chrysogenum* can parasitize *Calanus* copepods, causing elevated mortality and reduced swimming behavior. If this activity occurs in nature, we believe that predator avoidance, vertical migrations, and overwintering success could be reduced as a result of fungal colonization, underscoring the high likelihood that fungi play a role in the survival of key zooplankton species in the Arctic during the polar night.

## Acknowledgments

B. Hassett, E. Ershova, and J. Berge are jointly funded by UiT - The Arctic university of Norway and the Tromsø Research Foundation under the project “Arctic Seasonal Ice Zone Ecology”, project number 01vm/h15. We wish to acknowledge Randi Olsen from UiT for her assistance with zooplankton semi-thin sectioning and imaging.

## Author contributions

All authors contributed to the writing of this manuscript and study design. Brandon Hassett conducted fungal incidence surveys, culturing, and sequencing of fungal isolates. BH and Elizaveta Ershova conducted mortality experiments. EE conducted behavioral study analysis and determined zooplankton stage, identification, and sex. Jørgen Berge obtained funding and led the polar night expedition, as well as assisted in the acquisition of zooplankton specimens.

